# Protein Frustration Reveals Active Sites in Co-Evolved GPCR:G Protein Complexes and in Engineered Targeted Degrader Complexes

**DOI:** 10.1101/2025.06.27.660602

**Authors:** Wenyuan Wei, Roland Del Mundo, Tianyi Yang, Elizaveta Mukhaleva, Indira R. Sivaraj, Veerabahu Shanmugasundaram, Sergio Branciamore, Andrei S Rodin, Sivaraj Sivaramakrishnan, Ning Ma, Nagarajan Vaidehi

## Abstract

The folded structure of a protein is understood to be an optimal energy state. However, previous studies have shown that certain amino acid residue positions that play a critical role in protein function are often in a suboptimal energy state or “frustrated”. Here, we leverage over 1200 three-dimensional structures of G protein-coupled receptors (GPCRs) to demonstrate that residues at the interface between GPCR and its ligand or G protein contain a higher density of frustrated residues compared to other structural regions in the receptor. Likewise, the Gα subunit of the trimeric G proteins shows multiple clusters of highly frustrated residues on its surface that overlap with their effector protein (Gβγ, RGS, Adenyl cyclase, Ric8) binding interfaces. Compared to the co-evolved GPCR:G protein complexes, engineered protein complexes, such as those facilitated by molecular degraders, show a much greater density of highly frustrated residues in the degrader interface. Our study highlights the use of protein frustration as an invaluable tool to evaluate both native protein-protein interfaces and design strategies to facilitate engineered protein complexes.

## Introduction

The folding of the three-dimensional structure of protein complexes is driven by an interplay of physical and chemical interactions to minimize its free energy. However, not all regions within a protein or its complex end up in a stable low-energy state; some regions are in a sub-optimal energy state or “frustrated.” Protein frustration refers to the presence of competing interactions that prevent these regions from achieving their optimal, in isolation, energy state, resulting in local energetic conflicts, resulting in local energetic conflicts ^1^. Rather than being detrimental, frustration often plays a crucial role in biological function. Regions of high frustration are frequently associated with functional hotspots—sites where proteins interact with other molecules such as ligands or other proteins, undergo conformational changes, or catalyze chemical reactions^2^. These regions tend to be evolutionarily conserved and are strategically positioned to maintain functional flexibility while ensuring overall structural stability.

In protein complexes, the study of frustration has provided insights into key functional sites, such as binding interfaces, allosteric sites, and regions critical for enzymatic activity. By mapping the frustration landscape of a protein or protein complex, previous studies have shown how such energetic compromises are made to accommodate specific functional requirements ^3^. This approach has been increasingly employed in structural biology and drug discovery, offering a powerful framework for understanding protein function and guiding the design of therapeutics ^4^.

G protein-coupled receptors (GPCRs) form the largest superfamily of membrane proteins in the human genome, and they are also the largest target family for drug design ^5^. They serve as critical mediators of cellular communication by transducing extracellular signals, such as hormones, neurotransmitters, and sensory stimuli, into intracellular responses ^6^. This versatility is achieved through their ability to interact with a wide range of ligands and activate downstream signaling pathways via trimeric G proteins and other effectors ^7^. Despite their central role in drug development, designing ligands with high specificity and efficacy for GPCRs remains challenging. Their highly conserved structural framework, particularly in the transmembrane domain, poses challenges for designing drugs that selectively modulate specific GPCR subtypes without off-target effects. The dynamic nature of GPCRs, their ability to adopt multiple conformations, and the diversity of their ligand-binding mechanisms add complexity to the task. Analysis of the available three-dimensional structures of GPCRs has shown that ligands bind to multiple binding sites other than the orthosteric binding site and modulate the receptor activity.

A critical step in rational drug design is the identification of protein and/or ligand binding sites on GPCRs. These binding sites can be orthosteric, where endogenous ligands bind, or allosteric, which provide alternative regulatory sites. Understanding and identification of these sites are essential for designing molecules that can either mimic or modulate the receptor’s natural activity.

Proteolysis-targeting chimeras (PROTACs) have emerged as a powerful approach in drug discovery, enabling targeted degradation of disease-related proteins. Unlike traditional small-molecule inhibitors, PROTACs utilize a bifunctional design to recruit an E3 ubiquitin ligase to a target protein, leading to its ubiquitination and subsequent degradation by the proteasome ^8^. This innovative mechanism offers the potential to tackle previously “undruggable” targets and provides sustained therapeutic effects by eliminating rather than merely inhibiting the protein. Despite their promise, designing effective PROTACs presents significant challenges. One major challenge lies in maintaining a delicate balance between binding affinity, target selectivity, and degradation efficiency, as even minor modifications can disrupt PROTAC functionality. These challenges highlight the intricacy of PROTAC design and underscore the importance of integrating structural biology and computational modeling. Recently, we have shown in multiple PROTAC ternary complexes that frustration of residues in the ternary complex interface correlates well with the degradation efficacy of the PROTACs ^9,10^. In this study, we have applied residue pair frustration to identify protein and ligand binding sites in GPCRs and compared them to PROTAC ternary complexes.

## Results

### A protocol to calculate average frustration per residue from multiple GPCR structures

In this work, we have calculated the frustration of all the residue pairs using FrustratometeR ^11^. We downloaded 1283 three-dimensional structures of GPCRs that comprise Class A (1007), Class B (169), Class C (80), Class D (5), and Class F (22) structures from GPCRdb.org as of August 15, 2024 (see Supplementary Data). This dataset includes 850 active-state GPCR structures bound to either G protein (680) or β-arrestin (12), and 410 inactive-state structures bound to antagonists (232) or inverse agonists (48). For each of these structures, we employed mutational frustration FrustratometeR (V0.1.0, https://github.com/proteinphysiologylab/frustratometeR) to calculate the frustration index of every residue pair (see Methods, Fig. S1). Briefly, in the FrustratometeR procedure, each amino acid pair is mutated *in silico* to all other amino acids and the energy distribution for all the mutated pairs (decoy pairs) is calculated with respect to the energy of the wild type pair. The wild type residue pair energy is subsequently converted into a z-score (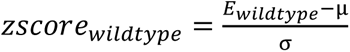, where μ and σ are the mean and standard deviation of the decoy energy distribution, respectively). The z-score is a measure of the wild type contact energy relative to all other amino acid substitutions at these residue positions, and hence an estimation as to whether a residue is in an energetically optimal or suboptimal state. The FrustratometeR toolkit is based on the coarse-grained forcefield AWESEM^32^. In each structure, each pair of residues from the output of FrustratometeR was classified as a highly frustrated (HF) pair if the z-score is <-1, neutrally frustrated (NF) (z-score is between 0.78 and -1), or minimally frustrated (MF) pair with z-score > 0.78 (Fig. 1A).

**Figure 1.**
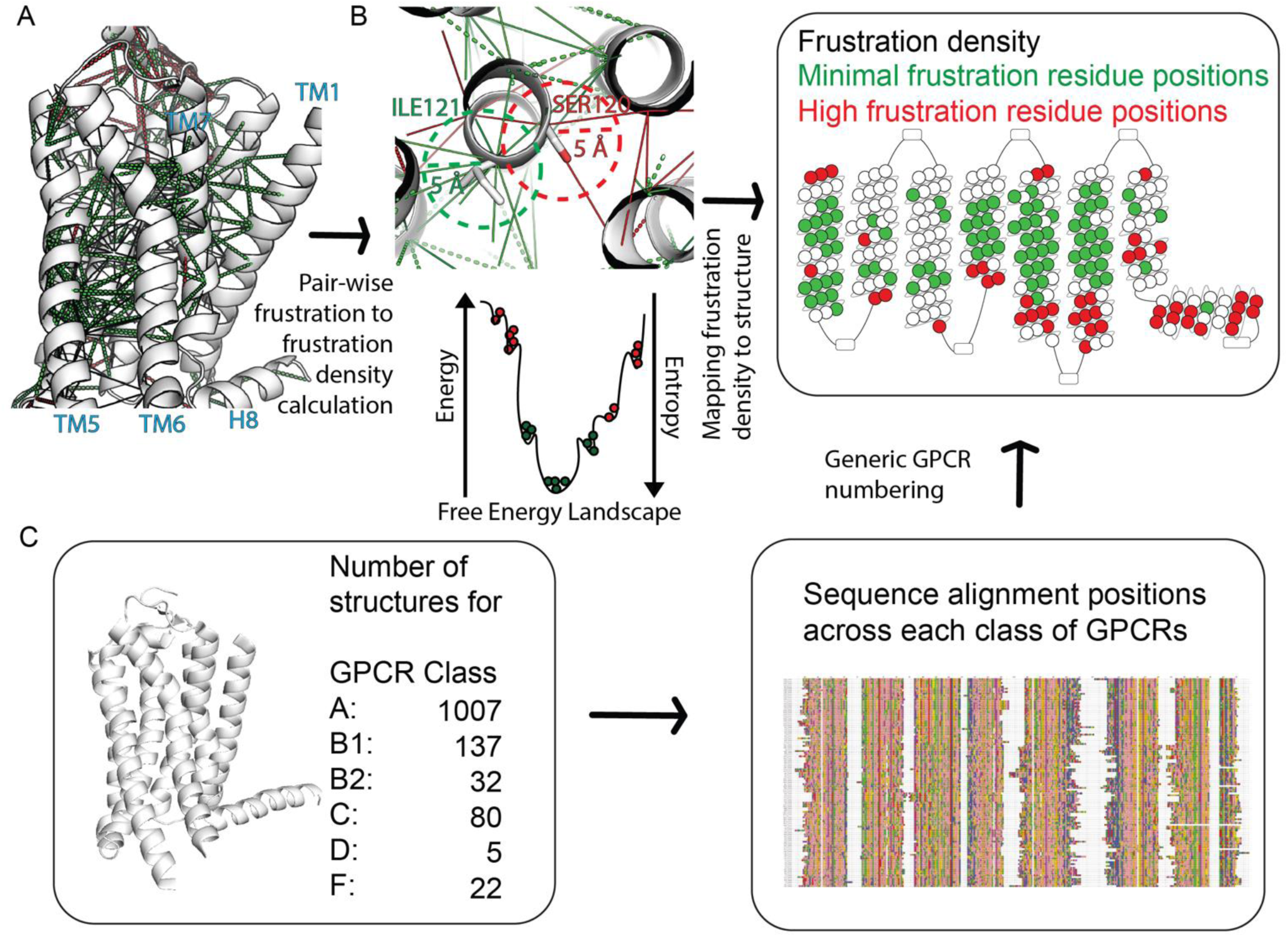
Workflow for calculating frustration. A) Residue pairwise frustration calculated using FrustratometeR (version 0.1.0, https://github.com/proteinphysiologylab/frustratometeR). Green dash and solid lines refer to minimally frustrated pairs, either water-mediated or direct-contacts, respectively. Red dash and solid lines refer to highly frustrated pairs either through water-mediated or direct-contacts, respectively. B) Procedure for conversion of the t pairwise frustration to single-residue frustration density. Residues within 5 Å of a given residue are considered for calculating frustration density. C) Workflow for calculating mean frustration density for each residue position in GPCR three-dimensional structures using the common residue numbering system derived from sequence alignments used for GPCRs.

We defined the frustration level of a protein-protein interface (PPI) as the percentage of HF or MF residue pairs in the interface to the total number of pairs in the interface 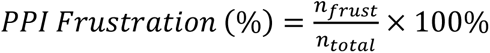, where 𝑛_𝑓𝑟𝑢𝑠𝑡_ is the number of highly or minimally frustrated interface pairs, and 𝑛_𝑡𝑜𝑡𝑎𝑙_is the total number of residue pairs in the interface. We then calculated the residue-based frustration as the sum of the frustration index of all the residue pairs within 5Å of the reference residue (Figure 1B). The percentages of HF, NF, and MF pairs within 5Å are calculated and assigned to the residue as its score in each of the three categories. We refer to these scores as HF density, NF density, and MF density, respectively. Residues in the top quartile of high frustration density are referred to as HF residues, while those in the top quartile of minimal frustration density are referred to as MF residues.

### Frustration is correlated to experimentally measured function and stability in Class A GPCRs

Figures 2A and 2B show HF and MF residues in class A GPCRs using the Ballesteros-Weinstein numbering system ^12^ (a total of 1007 structures, including both active and inactive state conformations) projected on the structure of adenosine 2A receptor A_2A_R (PDB ID 5G53) (Figs. 2A, 2B). Notably, we observed a high density of HF residues at the GPCR-G protein interface (red spheres in Fig. 2A), including the intracellular loops, whereas most of the MF residues were in the transmembrane helices (green spheres in Fig. 2B), rather than extracellular or intracellular loop regions. These observations suggest that HF residues may be linked to GPCR activity, given the importance of the GPCR-G protein interaction site for receptor function, while MF residues may support GPCR stability through the structural integrity of the transmembrane region. To find experimental support for this observation, we used the Receiver Operating Characteristic (ROC) analysis to check how well the HF density value of each residue retrieves the experimental positives.

**Figure 2.**
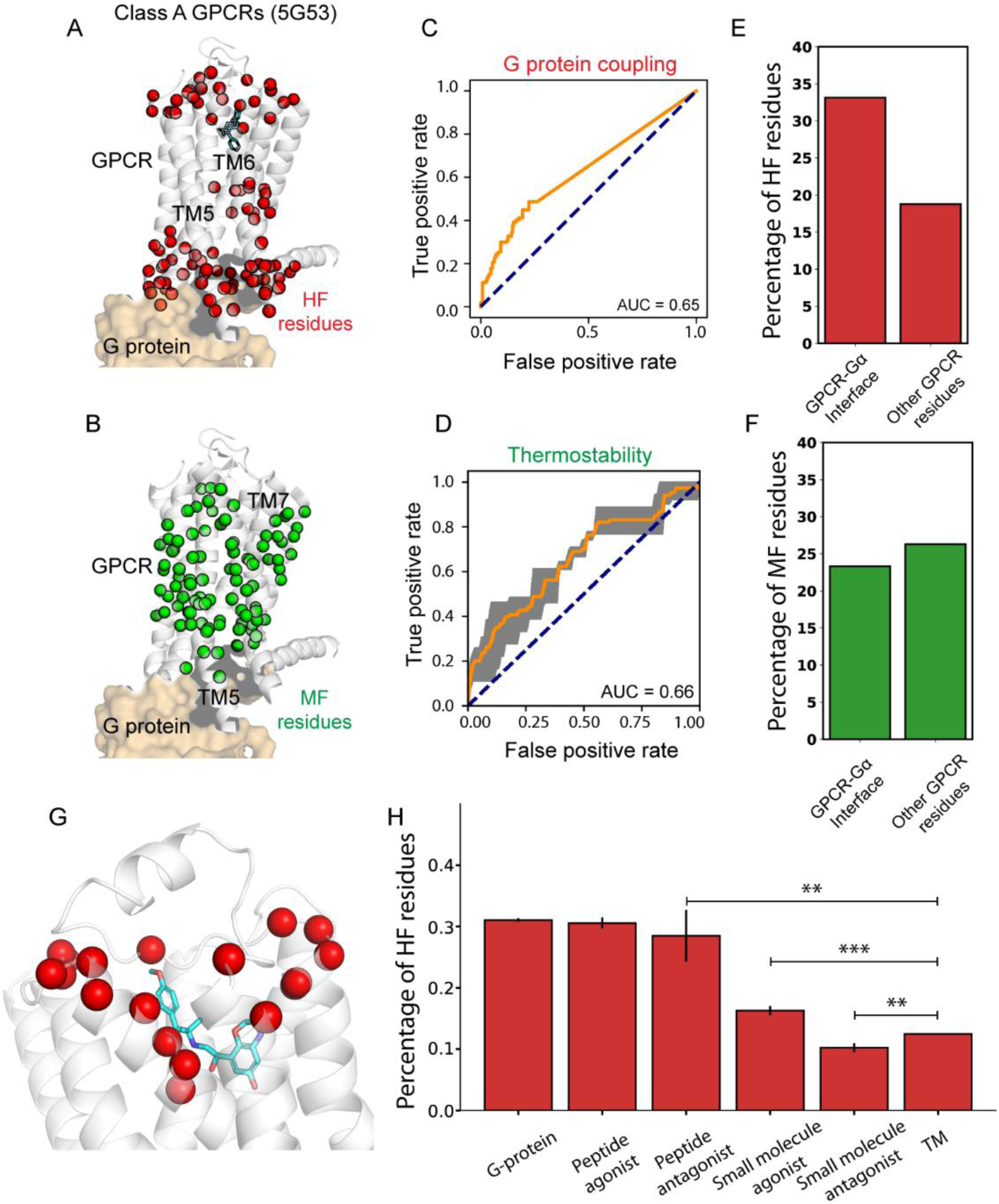
Frustration is correlated to the function and stability of Class A GPCRs. A, B) The HF residues (A) and MF residues (B) by their BW positions in class A GPCR (using 5G53 as template for displaying). C) The Receiver Operating Characteristic (ROC) comparing HF residues in AT1R and functional critical residues identified from Alanine scanning yields an Area Under the Curve (AUC) of 0.65. D) The ROC compares MF residues and thermostable residue mutations from Alanine scanning of adenosine A2A receptor (A2AR), (1-adrenergic receptor ((1AR), and neurotensin receptor (NTS1R), yielding an average AUC of 0.66. The average in ROC curve is shown as an orange line while the standard deviation of the ROC curve is shown as a gray shade. E, F) Percentage of HF residues (E) and MF residues (F) formed with each selected residue compared to GPCR-Gα interface residues and all other GPCR residues. G) HF residues around the ligand binding site among Adrenoceptor family (PDB ID: 8JJL). H) Percentage of HF residues in binding interfaces of G protein, peptide ligands, and small molecule ligands, regarding that of the TM region.

We used the experimental measurements of the change in the receptor coupling to the G protein upon mutation of each amino acid in the Angiotensin II receptor 1 (AT1R) to alanine. Briefly, the functional responses of each AT1R mutant for Gαq coupling were measured with the protein kinase C (PKC)–BRET biosensor ^13^. The dose-dependent responses to angiotensin II stimulation showed the level of change in both EC50 and Emax of angiotensin II in each AT1R mutant ^13^. The ROC analysis shown in Fig. 2C was performed using the frustration density calculated for each residue in AT1R and the measured difference in the receptor activity RA(Emax/EC50) between the mutant and the wild type AT1R. This analysis yielded an Area Under the Curve (AUC) of 0.65 (Fig. 2C), showing that HF density calculation of each residue from a single crystal structure recapitulates the receptor G protein coupling sensitivity better than random. Similarly, we used the MF density of each residue to check how well they retrieve the thermostability data from experiments. The thermostability data for the inactive adenosine receptor (A_2A_R), active β1-adrenergic receptor (β1AR), and active neurotensin receptor 1 (NTSR1) were taken from previously published data ^14^. The experimental thermostability measurements were performed on antagonist cyanopindolol bound turkey β1AR ^15^, agonist 5’-*N*-ethylcarboxamidoadenosine (NECA) bound adenosine receptor A_2A_R ^16^, antagonist ZM-241385 bound A_2A_R ^17^, and agonist NTS1 bound rat neurotensin receptor NTSR1 ^18^. Briefly, the experimental thermostability was measured by doing an alanine scanning mutation of each residue in each of these receptors and heating the detergent-solubilized mutant to an elevated temperature (∼28-32°C), cooling to 0°C, and the amount of correctly folded receptor was determined by a radioligand (either an agonist or antagonist or both) binding assay. In that work by Tate and colleagues, the thermostability of the wild type receptor was taken as 100%, and any mutants that showed 100% or higher ligand binding compared to the wild-type receptor were defined as thermostable. The ROC analysis of this data yielded AUC of 0.66 (Fig. 2D), which shows that the minimally frustrated residues are often important for structural stability of the GPCR. Taken together, these results supported our hypothesis that HF residues are likely associated with GPCR function, while MF residues contribute to GPCR stability. It should be noted that many of these residues contribute to both stability and function, but the frustration density shows the extent of leaning towards either stability or function.

We further investigated the location of the structural regions of the HF residues. Four distinct HF clusters of residues emerged: 1) the GPCR:G protein interface, 2) helix 8 (H8), 3) the well-characterized sodium binding site ^19^, and 4) three HF residues within the extracellular TM region, located near the orthosteric ligand binding site. The GPCR:G protein interface is well-established as critical for G protein coupling and hence the GPCR activity. Taking a closer examination, we observed that the bottom of transmembrane (TM) helices TM5 and TM6 within this interface contain a higher number of HF density residues compared to other TM helices. Notably, the intracellular (IC) region of TM5 and TM6 connects to intracellular loop 3 (ICL3), which, as shown in previous studies ^15,20^, plays a key role in GPCR activity and G-protein selectivity. This observation supported our hypothesis that HF residues are linked to GPCR activity. Residues in ICL2 were also highly frustrated (Fig. 2A). Due to the higher sequence diversity and difference in length in the loop regions, ICL1, ICL2, and ICL3 lack sufficient common residue positions for a systematic frustration analysis across all GPCRs. Therefore, we assessed each three-dimensional structure individually to evaluate the frustration differences between loop and TM regions (see method section for details. Fig. S1). Comparisons revealed that loop residues are generally more frustrated than TM residues, with a higher percentage of HF residues and a lower percentage of MF residues (Fig S2A, S2B). ICL2 is known to undergo a disorder-to-order structural transition during GPCR activation ^21^, so the high percentage of HF residues observed in ICL2 indeed correlates with GPCR activity. Recent studies have shown that helix 8 is critical for GPCR activity and G protein selectivity ^22–24^, while the sodium binding site is a well-known activation motif in class A GPCRs ^19^. The high percentage of HF residues observed in both regions (Fig. 2A) further supports our hypothesis.

Our study thus far suggested that frustration can serve as a metric for identifying functionally critical residues in GPCRs. We observed that the GPCR-G protein interface contains more HF residues compared to other regions of the GPCR (Fig. 2E, 2F). In contrast to the GPCR:G protein interface, the orthosteric ligand binding site showed fewer HF residues in the extracellular region. The orthosteric ligands in class A GPCRs bind in different structural locations, and therefore the number of common residue positions that we could average across class A GPCRs was very low. This variability could explain why we observed fewer HF residues around the orthosteric ligand binding site. Indeed, when analyzing only the adrenoceptor family of GPCRs (Fig. 2G), we observed a higher number of HF residues around the ligand binding site. To overcome the variability in ligand size, we calculated the percentage of HF residues in the binding interface of orthosteric ligands, categorized by their sizes and modalities (Fig. 2H). The TM residues that are not part of any binding interface were used as a control and labeled “TM”. While G-protein interface and peptide ligand interfaces all have higher percentage of HF residues compared to TM residues, signifying higher frustration in these interfaces, such cannot be easily claimed for small molecule ligands. In fact, small molecule antagonist binding interface is less frustrated than TM.

### Evolutionary properties of highly and minimally frustrated residues in GPCRs

If frustration is a functionally significant property, it is likely to be subject to evolutionary constraints. To investigate this, we assessed whether frustration correlates with sequence conservation and coevolution across Class A GPCRs. Using multiple sequence alignment (MSA) of 287 endogenous (without olfactory receptors) human Class A GPCRs, we calculated the conservation score (Cᵢ) based on Von Neumann entropy (S_i_) (see Methods). Specifically, for each alignment position, we constructed a density matrix ρ from the relative frequencies of amino acids, incorporating their chemical similarities through the BLOSUM50 similarity matrix. Because the maximum possible entropy in this setup is 1, the conservation score at each position is 𝐶_𝑖_ = 1 − 𝑆_𝑖_ . Applying this methodology to GPCR MSA revealed a substantial difference in average conservation between minimally and highly frustrated residues (Figs. 3A, B). MF residues were approximately 40% conserved, in contrast to only ∼20% for HF residues. This twofold increase in conservation for MF residues implies stronger evolutionary constraints.

**Figure 3.**
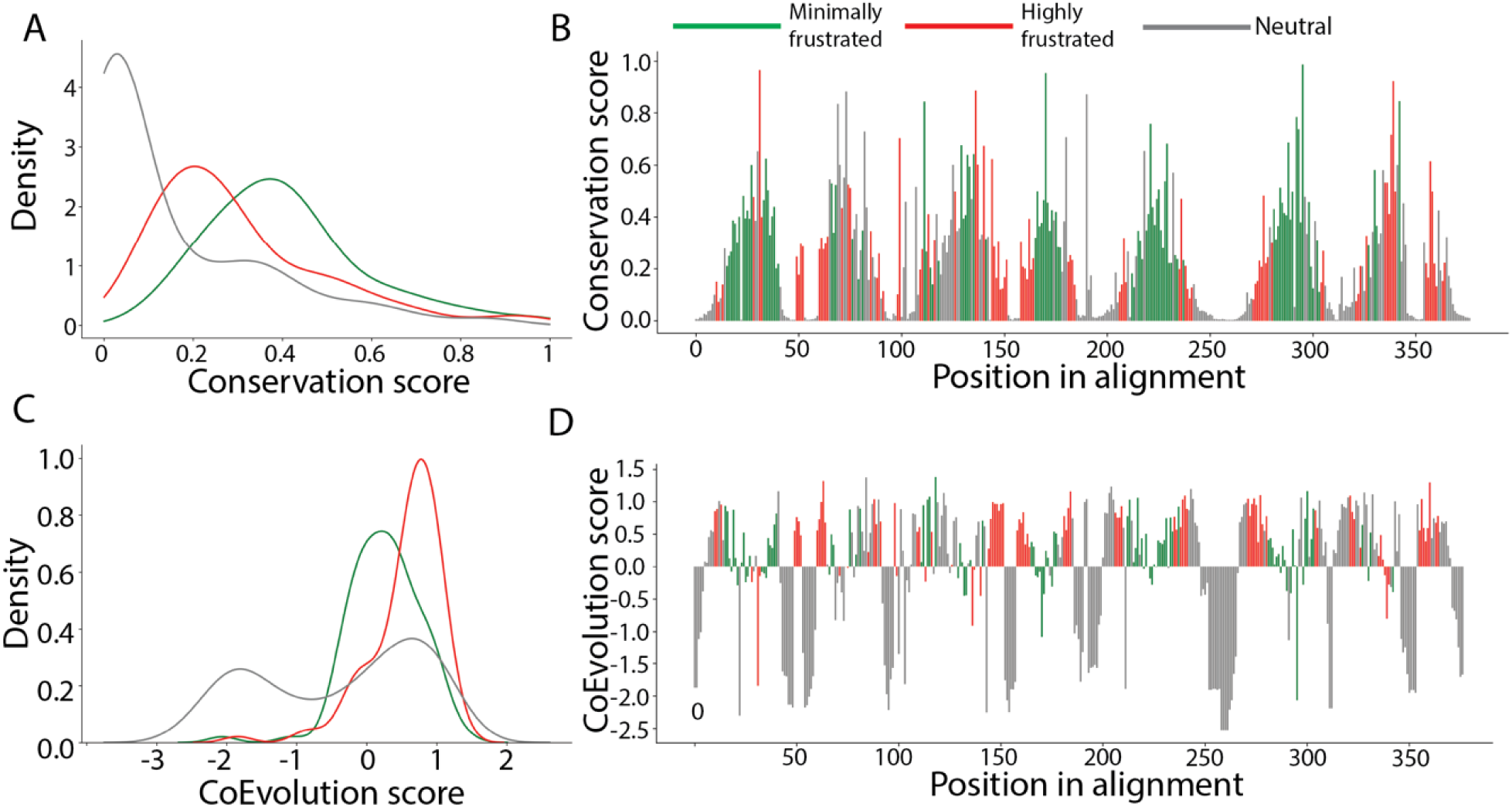
Sequence conservation and frustration. A. Population density plot of the conservation score of highly/neutral/minimally frustrated residues. B. Mapping of the conservation score for all residues in Class A GPCR alignment. Sets of minimally and highly frustrated residues are highlighted (green and red) and show a higher overall conservation score compared to neutral residues (grey). C. Population density plot of the coevolution score of highly/neutral/minimally frustrated residues. D. Mapping of the coevolution score for all residues in Class A GPCR alignment. Sets of minimally and highly frustrated residues are highlighted (green and red) and show a higher overall coevolution score compared to neutral residues (grey).

Next, we calculated the mutual information-based coevolution scores. Mutual information between two positions in the alignment measures how much knowing the amino acid at one position in the alignment reduces uncertainty about the amino acid at another position. We calculated the coevolution score for all positions in the same MSA (see Methods section) Figs. 3C, D). HF residues have coevolution scores one standard deviation above average. In contrast, MF residues clustered around average coevolution score values. These findings point to a compensatory or interdependent relationship for HF residues, whereas the elevated evolutionary stability of MF residues (high conservation, lower coevolution) suggests they act as structural “anchors.”

### Other classes of GPCR show a similar signature of frustration

We extended our study to include class B1 (137 structures, both active and inactive conformations, using the Wootten residue numbering scheme for this class ^25^), class B2 (32 structures, Wootten numbering scheme ^25^), class C (80 structures, Pin numbering scheme ^26^), class D (5 structures, Fungal numbering scheme ^27^), and class F (22 structures, Wang numbering scheme ^28^) GPCRs. The frustration patterns across these GPCR classes showed trends similar to those observed in class A GPCRs (Fig. 4).

**Figure 4.**
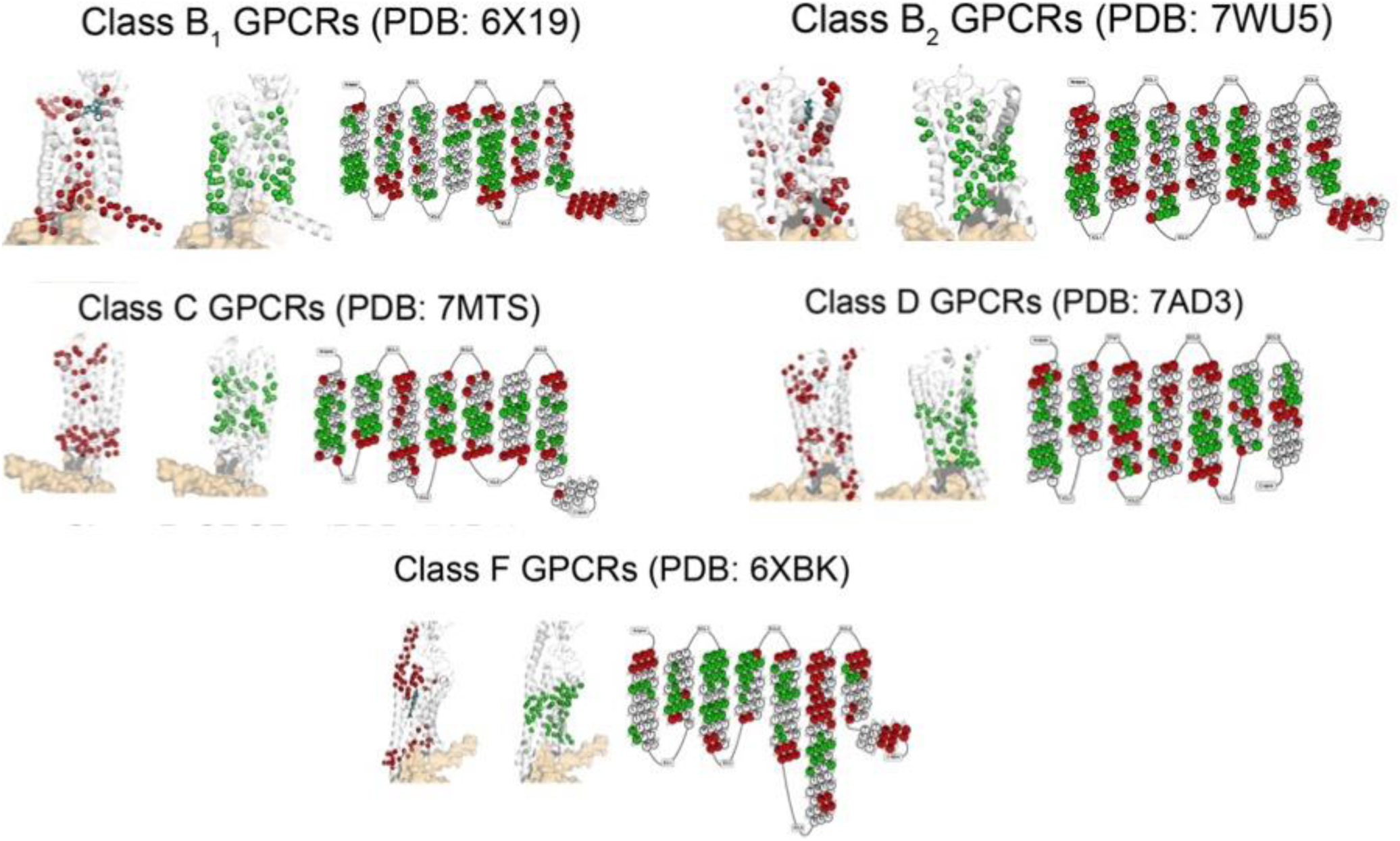
Location of highly and minimally frustrated residues in different GPCR classes. Frustration profiles were aligned using generic GPCR numbering for each class: Class B1 (Wootten), Class B2 (Wootten), Class C (Pin), and Class F (Wang), and averaged across available structures. HF and MF residues were mapped on representative structures Class B1: 6X19, Class B2: 7WU5, Class C: 7MTS, Class D: 7AD3, Class F: 6XBK. Native ligands are shown as cyan sticks to highlight the position of the orthosteric ligand binding site. On the right, snake plots show HF and MF residues, mapped on representative plots for each class: GLP-1 receptor (Class B1), ADGRF1 (Class B2), mGlu2 receptor (Class C), STE2 receptor (Class D), and SMO receptor (Class F).

The G protein binding sites and helix 8 exhibit high frustration, indicating the functional importance of both regions across all GPCR classes. The orthosteric ligand binding sites on the extracellular (EC) side are also highly frustrated, except in class C GPCRs, where their orthosteric binding sites are in the EC domain rather than within the TM bundles. The EC domains and loop regions also exhibit high frustration, reflecting their critical roles in receptor function (Fig. S3). For example, the EC domain of the GLP-1 receptor, which facilitates GLP-1 binding, displays high frustration ^29^. Similarly, the EC domain of the metabotropic glutamate mGlu2 receptor, known as the Venus flytrap domain, is highly frustrated and essential for glutamate binding and receptor dimerization ^30,31^. In class D, Ste2 receptor, both the dimerization interface and ligand-binding sites are highly frustrated ^32^. Likewise, the EC domain of the SMO receptor, which includes a cysteine-rich domain, exhibits high frustration at binding sites for oxysterols, highlighting its role in the Sonic Hedgehog signaling pathway ^33^. Finally, the central region of the TM bundle shows lower frustration, as indicated by a higher proportion of minimally frustrated residues.

### The protein-protein interface on the Gα subunit is highly frustrated

The trimeric G protein has three subunits, namely the Gα and Gβγ. The Gα subunit has multiple subtypes. In this section, we analyze the frustration residue pairs in the Gα subunit of the trimeric G proteins. We begin by analyzing the GPCR:Gα interface. Like GPCRs, G proteins have a Common G alpha Numbering (CGN) system ^34^, which allows us to align different Gα subtypes (G_αS_, G_αI/O_, G_αQ/11_, and G_α12/13_ for which three dimensional structures are available) and calculate the average frustration for each aligned residue position. By combining CGN numbering for Gα protein with residue numbering for GPCRs, we have identified the frustration residue pairs along the GPCR-G protein interface (Fig. 5A). We observed high numbers of both HF and MF pairs between TM3, TM5, and ICL2 of the GPCR and the H5 helix of the G protein (Fig. 5B).

**Figure 5.**
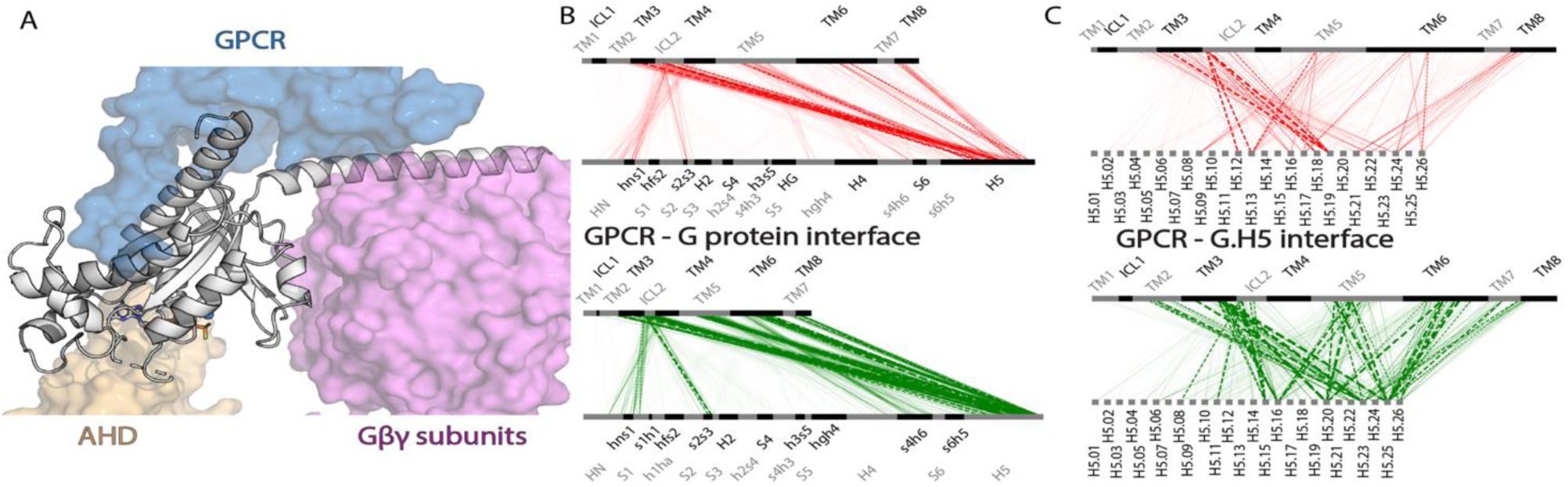
Residue pairwise frustration in the GPCR-Gα interface. A) A structural overview of the interface between GPCR and Gα. AHD is the all helical domain of the Gα subunit. The Ras domain of Gα is shown in a grey cartoon. The Gβγ subunits are shown in a pink surface. B) HF (top) and MF (bottom) pairs between GPCR and Gα across all structural regions. C) HF (top) and MF (bottom) pairs between GPCR and the H5 helix of Gα. The thickness of the line reflects the percentage of said pairs occurring in all structures.

Since the H5 helix is known to be critical for Ga coupling to GPCR, we examined the residues forming HF and MF pairs on the 27 amino acids of the H5 helix (Fig. 5C). The HF and MF residues on H5 helix in the Ga protein are distinct: the middle section of H5 (H5.09-H5.19) contains the HF pairs, whereas the C terminus of H5 (H5.20-H5.26) contains the MF pairs. This observation suggests that the C terminus tip of H5 helix is likely crucial for providing stability to the GPCR:G protein complex. The middle section of the H5 contributes to selectivity of the Gα couplings. These observations have experimental support from previous studies ^35–40^.

The Gα subunit of the G protein has four major subtypes (G_αS_, G_αI/O_, G_αQ/11_, and G_α12/13_) and samples three conformational states: GDP-bound trimer state, GPCR-bound complex state, and GTP-bound monomeric state. We downloaded 637 three-dimensional structures of Gα proteins in monomeric and effector-bound (84), trimeric (3), and GPCR-bound (550) conformations as of August 15^th^, 2024. We first calculated the frustration density of residues in the Ga subunit in the GDP-bound trimer state. Surprisingly, in the GDP-bound trimer state, which is the “off” state, the H5 helix does not contain any HF residues, even though we observed that residues on H5 are highly frustrated when coupled to GPCRs. Most of the HF residues are found at the interface between the Ras domain and AHD, with additional HF residues located in the Gα-Gβγ interface and other regions along the Gα-GPCR interface outside of H5 (Fig. 6A). The Ras-AHD interface is crucial for nucleotide binding, and it has been shown that the AHD domain demonstrates dynamic motion with respect to the Ras domain during GPCR-dependent activation to enable the nucleotide release and exchange ^41,42^. These well-established concepts indicate that the Ras-AHD interface is critical for G protein function, so the high frustration observed there aligns with functionality. Similarly, the Gα-Gβγ interface is essential for G protein activity due to the requirement for Gα-Gβγ association and dissociation, agreeing with our observation of the number of HF residues in that region.

**Figure 6.**
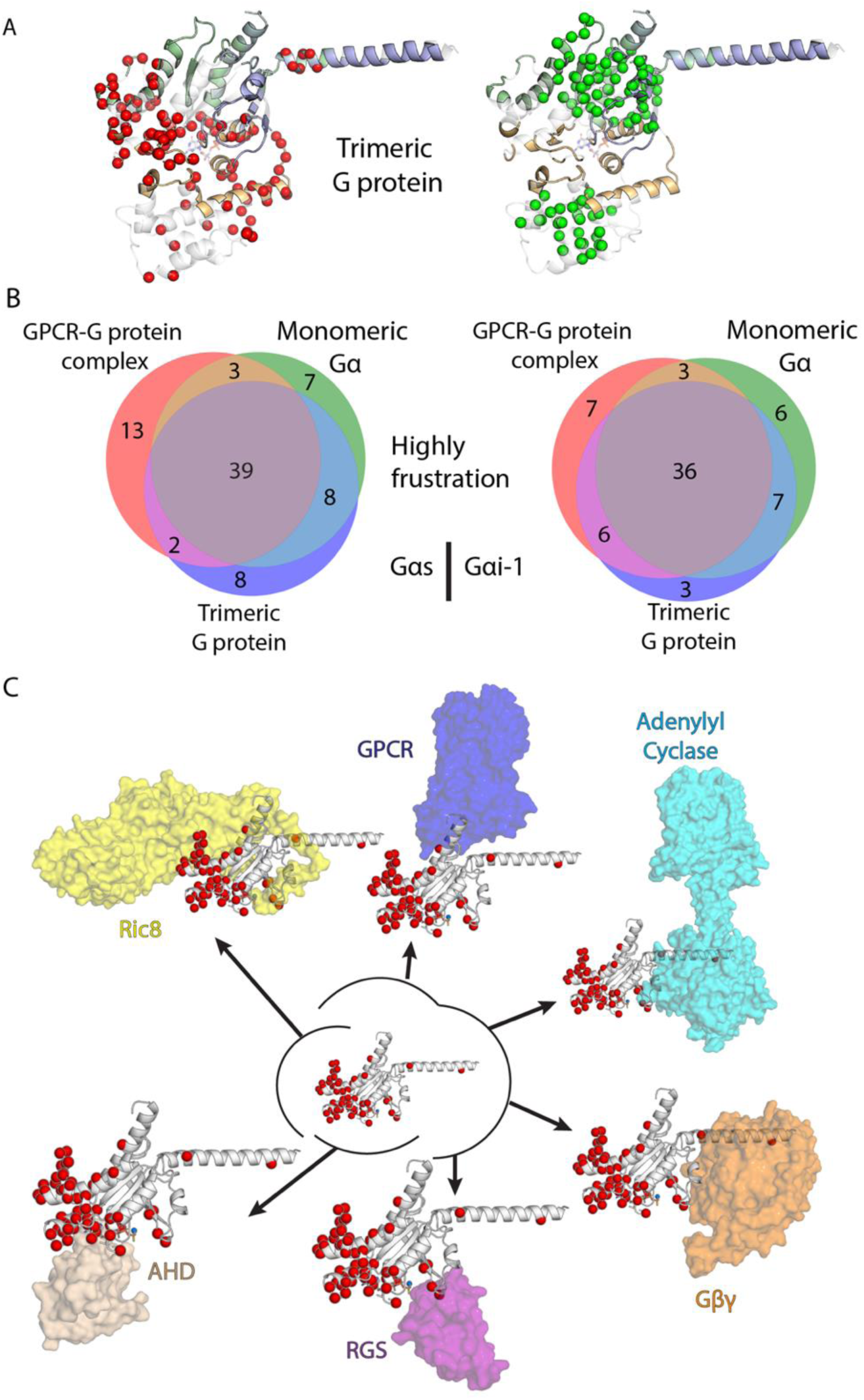
Frustration profiles of Gα in Gβγ-bound trimer and high frustration interface characterization of Gα in the three states. A) High frustration (red spheres - left) and minimal frustration (green spheres - right) residues in Gα subunit in GDP-bound GαGβγ-bound trimer state. B) Venn diagram of the HFRs in three conformational states (GDP-bound trimeric state, GPCR-bound intermediate state, and GTP-bound Gα monomeric state) illustrating the common and distinct HFRs in the three states in Gα_S_ (left) and Gα_I-1_ (right). C) Alignment of Gα Ras domain with protein effectors of Gα; frustration profile from the GPCR-bound trimer state is shown.

We compared the similarities in distributions of HF and MF residues among the GDP-bound trimer, GPCR-bound complex, and GTP-bound monomeric states in two Ga subtypes, namely G_αS_ and G_αI_ (Figs. 6A, 6B, and S4A). Interestingly, 39 of the HF residues in G_αS_ and 40 HF residues in G_αI_ are common to all three conformational states (Fig. 6B). As shown in Fig. 6C, the highly frustrated residues are located on different protein (such as GPCR, Gβγ, RGS, Ric8, adenyl cyclase) binding interfaces in the Gα subunit, indicating that protein binding interfaces are frustrated. The information on the protein binding interfaces on the Gα subunit was derived from the three-dimensional structures available in the PDB on August 15^th^, 2024.

We then examined the distribution of the HF residues that are common to G_αS_ and G_αI_ in the Ras domain structures. We observed a high number of common HF residues in HG-hgh4-H4 region in both G_αS_ and G_αI_ (Fig. S4C). This region is the hinge of Ras-AHD domain movement and part of the Ras-AHD interface and therefore possesses significant functional importance. The same analysis was also done for the shared MF residues. Similarly, many MF residues were shared among the three states (Figure S4B), and they are located mostly on structured regions (S1, S3, S4, S5, and H3) of the Ras domain of Gα subunit (Figure S4D). This implies their importance in structural stability. When comparing frustration in G proteins and GPCRs, a significant difference emerges: the distributions of HF and MF residues were less clearly separated in G proteins. In GPCRs, MF residues were primarily located in the TM bundle, whereas in G proteins, HF and MF residues showed a notable overlap in different structural regions (Fig. S4A). One possible explanation for this difference is that G proteins have multiple functions and interact with various downstream proteins. As a result, much of the G protein surface is likely functionally relevant in different contexts, leading to a generally higher level of frustration: we noticed that the Ras domain is more frustrated than the AHD, consistent with its role as the primary functional region of the Gα subunit^43^.

### Frustration difference in the interface of co-evolved protein complexes compared to engineered protein degrader ternary complexes

So far, we have discussed frustration in GPCR-G protein complexes, which are endogenous and co-evolved protein complexes. However, there are also many protein complexes that are engineered. Targeted protein degradation is an area of research where small molecules are designed to form complexes between the protein of interest (POI) and ubiquitin E3 ligase, to ubiquitinate POI and degrade it. There are two major types of degrader molecules: PROTACs (PROteolysis Targeting Chimeras), and Molecular Glues. PROTACs are composed of two binding small molecules—one targeting the POI and the other binding to E3 ligase, and these two moieties are connected by a linker. This modular design enables PROTACs to mediate interactions between proteins that may not naturally bind, creating what we call the “engineered protein-protein interfaces”. Molecular Glues function by stabilizing transient or weak interactions between two proteins, effectively enhancing their natural affinity. Unlike PROTACs, which rely on a bifunctional linker to bridge the POI and E3 ligase, molecular glues promote degradation by inducing conformational changes that facilitate direct recruitment of the POI to the E3 ligase ^44,45^. Designing PROTACs and molecular glues poses a challenge due to the dynamic nature of the ternary protein complexes formed. Structure-based design principles governing these engineered complexes would greatly aid the design of these degraders. Here we studied the frustration patterns of residues in the protein-protein interface for 36 PROTAC-mediated ternary complex structures and 50 molecular glue-mediated complexes (date of download: Jun.13^th^ 2024) and compared them to frustration patterns observed in GPCR:G protein complexes. The three-dimensional structures of molecular glue degrader complexes included the E3 ligases (cereblon, VHL and DDB1) and the POIs (IKZF1/3 and GSPT1) (details in the Methods section). The POIs in the structures for PROTAC-based ternary complexes include BRD4, WDR5, SMARCA2, SMARCA4, KRAS, FAK, FKBP51, and BCL-XL. VHL (von Hippel-Lindau disease tumor suppressor) is the E3 ligase in most of the PROTAC-based ternary complex structures (29 out of 36), and the rest of the structures contain cereblon. We calculated the average residue-based frustration values for all the residues (as described before and in Methods) in 29 VHL-POI ternary complexes (see Methods) and projected the frustration in the structure (Fig. 7A). Most of the HF residues are located in the PROTAC binding site in the VHL-POI interface. The VHL-Elongin C (blue surface in Fig. 7A) interface in the E3 ligase complex that have naturally evolved to form a complex also shows a higher density of HF residues than GPCR complexes but lower than PROTAC ternary complexes. These findings align with our previous conclusion that HF residues are concentrated in protein-protein interfaces. A similar pattern was observed in the cereblon-based PROTAC ternary complexes as well as DDB1 molecular glue systems (Fig. S5A, B)

**Figure 7.**
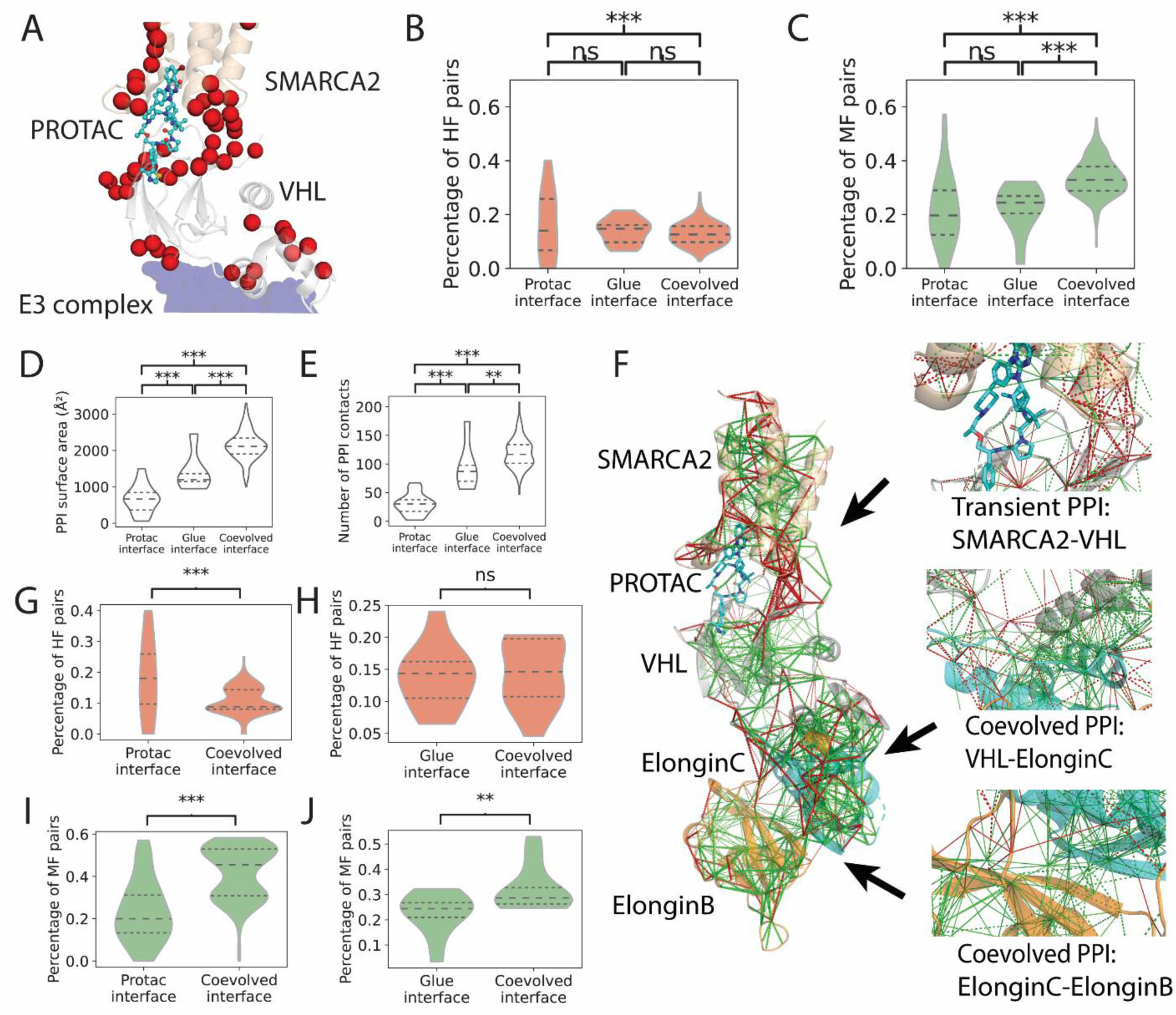
Application of frustration in target protein degrader (TPD) design. The data shown for co-evolved complexes include GPCR:G protein complexes and protein interfaces in E3 complex, such as VHL:Elongin C and Elongin C: Elongin B. A) An example of highly frustrated residues mapped on VHL-TPD-SMARCA2 ternary complex (PDB id: 7Z76). The TPD molecule is colored cyan. The SMARCA2 is colored light brown. The VHL E3 ligase is colored gray. Elongin-C (component of E3 ligase) is colored dark blue. The top quartile of highly frustrated residues is shown in red spheres. B, C) High and minimal frustration level of residues in the protein-protein interface for PROTAC-mediated interface, molecular glue-mediated interface (excluding DDB1-related complexes), and co-evolved interface (GPCR-G protein interface). D) PPI surface area. E) Number of PPI contacting residue pairs. F) Pair-wise frustration projected onto SMARCA2-E2 complex. One engineered interface (SMARCA2-VHL) and two co-evolved interfaces (VHL-ElonginC, and ElonginC-ElonginB) are shown in insets. The pair-wise frustration shows that engineered interface is more highly frustrated (more red dash lines) compared to co-evolved interface. G, H) High PPI frustration level for PROTAC and molecular glue complexes. The co-evolved interface refers to the PPI between components that naturally associate in E3 ligase complex, such as the CRBN–DDB1 and VHL–Elongin C interfaces. I, J) Minimal PPI frustration level for PROTAC and molecular glue complexes. The co-evolved interface refers to the PPI between components that naturally associate in E3 ligase complex, such as the CRBN–DDB1 interface.

We compared the percentage of HF residue pairs in the engineered degrader complex interface to the co-evolved GPCR:G protein complex interface. Our analysis revealed that the PPIs of engineered degrader complexes contain a higher percentage of HF pairs compared to GPCR:G protein complexes (Fig. 7B). Notably, PROTAC-mediated PPIs show a higher percentage of HF residue pairs compared to molecular glue-mediated interfaces (Fig. 7B). Conversely, co-evolved GPCR:G protein interface shows a higher percentage of MF pairs than the degrader complexes (Fig. 7C), with a similar difference observed between PROTAC- and molecular glue-mediated PPIs. The frustration analysis for molecular glue mediated-DDB1-CDK12 interfaces is shown in (Fig.S6 A, B). These findings indicate that engineered PPIs are more frustrated overall than co-evolved PPIs. As shown in Fig. 7D, the GPCR:G protein complexes demonstrate a larger buried surface area in the interface (1000–3000 Å²) than PROTAC ternary complexes (50–1500 Å²) and molecular glue-mediated interfaces (950–2500 Å², excluding DDB1-related complexes). As a secondary validation, we analyzed the number of contacts in each PPI (Fig. 7E) and found that GPCR:G protein interfaces contain more contacts than PROTAC-mediated PPIs and molecular glue-mediated interfaces (excluding DDB1-related complexes). Interestingly, the molecular glue– mediated DDB1:CDK12 interfaces are distinct from other degrader-mediated interfaces. Their buried surface area (BSA) and number of contacts are even greater than those observed in GPCR:G protein interfaces (Fig.S6 C, D). Both measures confirm that molecular glue- and PROTAC-mediated interfaces are distinct in their structural characteristics, despite both being more frustrated than GPCR-G protein interfaces.

We also compared frustration residue levels within the same ternary complex by examining both engineered and co-evolved interfaces in the POI-E2/E3 ligase complex. Using the SMARCA2-PROTAC-VHL-ElonginC-ElonginB complex as an example, we analyzed one engineered interface (SMARCA2-VHL) and two naturally co-evolved interfaces (VHL-ElonginC and ElonginC-ElonginB). The results show that the engineered interface exhibits higher number of HF pairs than both co-evolved interfaces (Fig. 7F), consistent with the observation on all the PROTAC (36 structures) and molecular glue (50 structures) complexes (Fig. 7G, H, I, J, Fig.S6 E, F). These findings confirm that the differences we observed are not specific to comparisons with GPCR:G protein interfaces but hold true across different systems, further supporting the distinct frustration patterns of engineered versus co-evolved protein-protein interactions.

### Application of frustration in identifying allosteric modulator sites in GPCRs

Allosteric modulators are sought after in GPCR drug design yet remain a challenge. These modulators bind to allosteric sites and either enhance (positive allosteric modulators, PAMs) or reduce (negative allosteric modulators, NAMs) the pharmacological properties of the orthosteric ligand. To investigate whether residue frustration density can be used to identify such sites, we used all available Class A GPCR structures containing PAMs or NAMs from GPCRdb.org and clustered the PAM and NAM binding sites based on the geometric center of the ligand heavy atoms. This analysis revealed six locations of the binding sites for PAMs and four for NAMs (Fig. S7A, S7B). We found that PAM binding sites contain a higher percentage of HF residues and NAM binding sites have a lower percentage of HF residues than the TM region, (Fig. 8A). The TM region is defined as the region in the transmembrane domain that does not contain these binding sites. This observation is similar to that of agonists and antagonists shown in Fig. 2H. Additionally, each PAM/NAM binding site contains several HF residues (Fig. S7A, S7B). We observed a similar trend for the PAMs and NAMs in other GPCR classes (Fig. S7C). This finding further supports the use of frustration as a tool for identifying potential allosteric ligand binding sites in GPCRs. Next, we analyzed how these allosteric modulator sites can be identified using the structure with no PAM bound to the receptor. Such information will be useful for designing PAMs for GPCRs.

**Figure 8.**
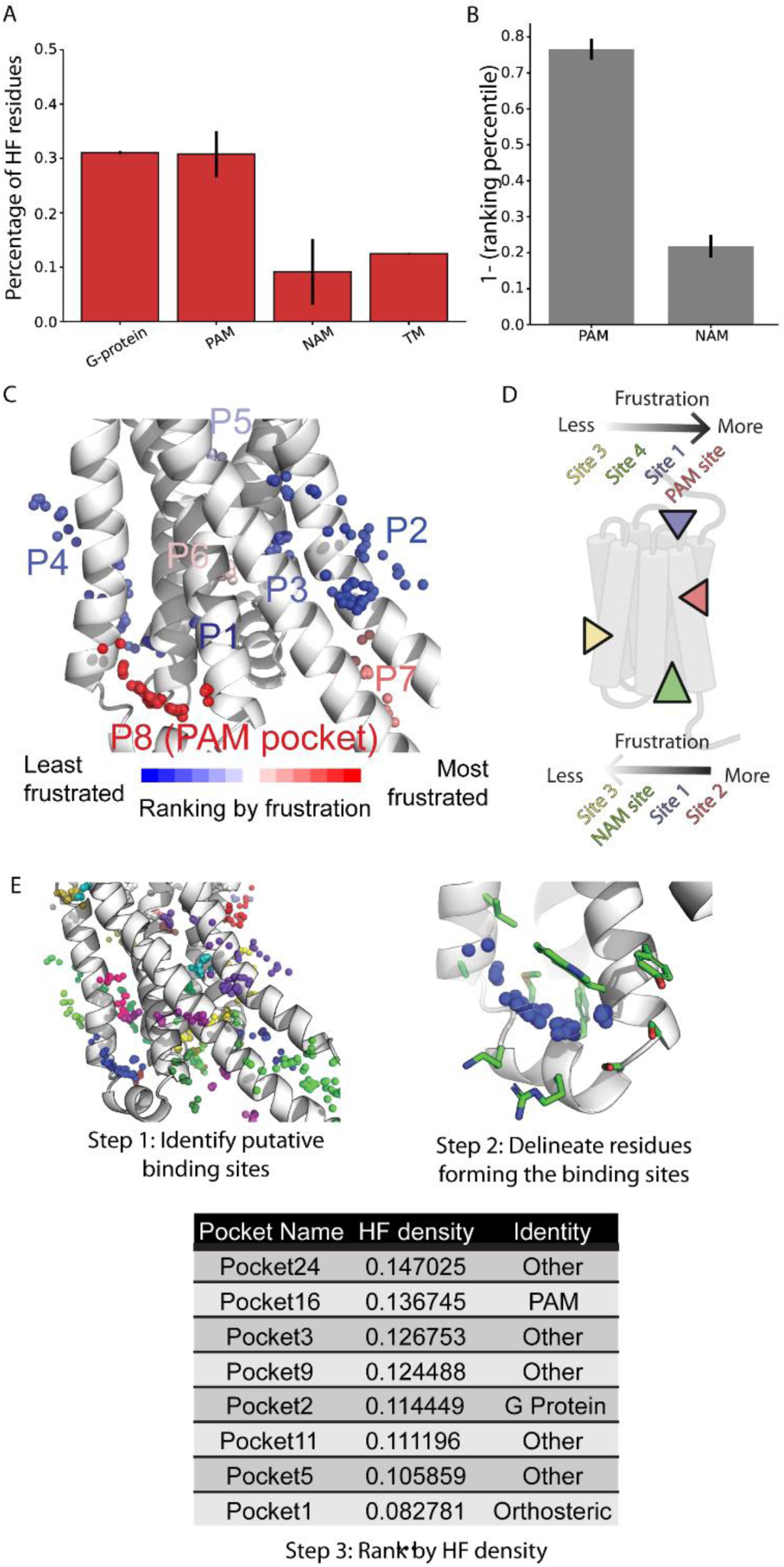
Predicting GPCR allosteric modulator binding sites using frustration. A) Percentage of HF residues in PAM and NAM binding sites. G-protein binding sites and TM regions are used as positive and negative controls, respectively. B) Ranking of true PAM and NAM binding sites (for all class A GPCRs where structures with PAM and NAM are available) among all putative binding pockets in the TM regions. The Y-axis values are 1 – quartile in which the binding site appears. C) In the D1 structure (PDB ID: 7JOZ), the known PAM binding site is the most frustrated among all putative binding sites in the TM regions. All putative binding sites discovered by fpocket are shown as spheres and colored by their mean HF density. D) A schematic figure showing how to apply frustration to identify true PAM/NAM binding sites.

As a case study, we compiled all Class A GPCRs with known PAM or NAM co-resolved in the structures. These receptors also have three-dimensional structures without the modulators. The GPCRs with known PAMs include human CB1R, FFAR1, D1R, GPBA, HCA2, M2AChR, M4AChR, MRGPRX1, TSH, β2AR, NTSR1, and rat NTSR1, while those with known NAMs include human C5a1R, CB1R, M1AChR, and β2AR. We collected all active-state structures without PAMs and all inactive-state structures without NAMs. We developed a protocol to identify a modulator binding site as enumerated below (Fig. 8D, E):

1. Identify potential ligand binding sites in each three-dimensional structure using the fpocket algorithm (https://doi.org/10.1186/1471-2105-10-168).
2. Calculate the mean HF density of all residues in a binding pocket for each putative ligand binding site.
3. Rank the putative binding sites for each structure from the most to the least frustrated.

We observed that the true PAM binding site ranks second by the average HF density in the case of β2AR. However, the correct PAM binding site falls within the top quartile of the average HF density distribution and the NAMs site falls within the last quartile of the average HF density (Fig. 8B, C). This suggests that frustration can be used in identifying PAM/NAM binding pockets. This suggests that frustration can be used in identifying PAM/NAM binding pockets, which, followed by Virtual Ligand Screening in these sites, could aid modulator design.

## Discussion

Previous studies on energetic frustration have shown that highly frustrated residues are frequently located at active sites in proteins. Here, we leverage the vast database of published GPCR structures to demonstrate that protein frustration can also identify sites of interactions with ligands and G protein heterotrimers. Interestingly, protein frustration reveals high frustration not only in the orthosteric ligand binding site but also in the allosteric sites, which are not sites of action of the endogenous ligands. Our study demonstrates that evolution might have optimized residues at protein-protein interfaces to be passivated by the coupling interaction. In parallel, we find that protein frustration is a helpful tool to facilitate interactions in engineered complexes to enhance the efficiency of molecular degraders that operate at these non-native interfaces. Hence, protein frustration serves as an accessible tool to evaluate the chemistry of engineered interfaces to facilitate complexation.

In this study, we computed residue-level frustration for over 1200 GPCRs and their complexes with the G proteins. Our analysis revealed that highly frustrated residues cluster predominantly at the G protein-coupling interface and ligand-binding sites of GPCRs. Our calculated results were validated experimentally using the mutations of these highly frustrated sites affect G protein coupling. This was established by correlating frustration scores with experimentally measured coupling strength using bioluminescence resonance energy transfer (BRET) assays. In contrast, minimally frustrated residues are predominantly located within the central regions of transmembrane helices. These residues contribute to receptor stability and align well with thermostability experimental data. Moreover, minimally frustrated residues tend to be more evolutionarily conserved than highly frustrated ones, whereas highly frustrated residues exhibit greater co-evolutionary values.

Our findings suggest that analyzing frustration from static structural snapshots can serve as a powerful approach to identify novel druggable allosteric sites in GPCRs. Interestingly, agonist binding sites show a higher density of highly frustrated residues compared to antagonist sites, potentially allowing greater flexibility to facilitate the conformational changes required for receptor activation.

Analysis of the Gα subunit of trimeric G proteins—regardless of subtype (G_αS_, G_αI/O_, G_αQ/11_, and G_α12/13_) or activation state (“on”/“off”)—revealed multiple clusters of highly frustrated surface residues. These clusters coincide with known interfaces for a range of effectors, including Gβγ, GPCRs, RGS proteins, Ric8, and adenylyl cyclase. Within the H5 helix—a critical structural element for GPCR coupling—we observed that minimally frustrated residues at the helix tip may stabilize GPCR binding, while highly frustrated residues mid-helix could act as a lever to facilitate nucleotide exchange by modulating the opening and closing of the binding site. This dynamic behavior has been previously described in molecular dynamics simulations ^42^.

Structure-based drug design has traditionally struggled with targeting molecular degraders due to the highly dynamic nature of their ternary complexes. Our frustration analysis of ternary complexes formed by both PROTACs and molecular glues with E3 ligases showed a greater density of highly frustrated residues at the interfaces involving the degrader, target protein, and E3 ligase in comparison to the GPCR:G protein interface. This suggests that such engineered complexes are less evolutionarily optimized and inherently more transient. In conclusion, our work underscores the value of understanding residue-specific energy landscapes in protein complexes. Such insights can inform novel strategies for designing therapeutics that target dynamic protein-protein interfaces.

## Materials and Methods

### Frustration calculation via FrustratometeR

FrustratometeR was downloaded and installed on the local cluster from GitHub repository (https://github.com/proteinphysiologylab/frustratometeR) in a Conda environment. Mutational frustration was calculated for each structure with a R submission script, with all other parameters set to default. The results included two files: [pdb].pdb_mutational, which is the pair-wise frustration data, and [pdb].pdb_mutationa_5adens, which is the frustration density of each residue within 5Å. Both were then parsed using in-house scripts for further analyses.

### GPCR structure information

A total of 1283 GPCR structures were collected from GPCRdb (https://gpcrdb.org/) for analysis, including 1007 Class A, 137 Class B1, 32 Class B2, 80 Class C, 5 Class D, and 22 Class F GPCRs. This dataset includes 850 active-state GPCR structures bound to either G protein (680) or arrestin (13), and 410 inactive-state structures bound to antagonists (232) or inverse agonists (48). Species and chain information was retrieved from GPCRdb (https://gpcrdb.org/structure/) using Structure API. PDB structure files were downloaded on August 15, 2024.

### G protein structure information

A total of 636 Gα structures were collected from GproteinDB (https://gproteindb.org/) for analysis, including 3 Gα-Gβγ trimer, 25 Gα monomer, 57 Gα-effector complex, and 551 GPCR-Gα-Gβγ complex. Structures that were labeled to have chimeric Gα were excluded. These structures are grouped based on Gα states into GPCR-Gα complex (551), Gα-Gβγ trimer (3), and Gα monomer (82). Species and chain information was retrieved from RCSB (https://www.rcsb.org/) using the REST API.

### Contact analysis

We employed “get_static_contacts.py” module from GetContacts (http://getcontacts.github.io/) for contact analysis. The output TSV file was further processed using in-house code to retrieve the residues lining up the interface. Briefly, given the chain ID of interest, the entity containing the chain was kept in each row; the residue number was extracted from the entities, from which a unique list was created.

### GPCR-Gα frustration pairs

Class-A-GPCR-Gα complex data were used in this analysis. Frustration pair information was parsed from [pdb].pdb_mutational files. Pairs made between GPCR and Gα that are classified as either highly or minimally frustrated were kept for analysis. Residues of each structure were first converted to BW and CGN for GPCR and Gα, respectively. A super set of contacts was collected and grouped by the contacts’ structural regions, according to BW and CGN. The plot was generated using the Python Matplotlib.pyplot package. The frequencies of each contact present were calculated and used as the width of lines.

### Per-residue frustration density, mean residue frustration density, and mean region frustration density calculation

The per-residue frustration density 𝜌*_i_*^𝑋^was computed as:

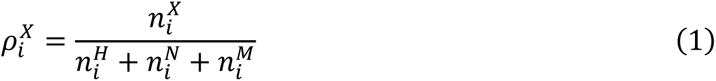

where each residue 𝑖 interacts with residues within a 5 Å sphere (𝑛*_i_*^𝑋^), generating frustrated pairs categorized as highly frustrated (𝑛*_i_*^𝐻^), neutrally frustrated (𝑛*_i_*^𝑁^), or minimally frustrated (𝑛*_i_*^𝑀^). The frustration type is denoted by 𝑋 ∈ {𝐻, 𝑁, 𝑀} . For GPCR per-residue frustration density, frustration profiles were aligned using generic GPCR numbering specific to each class: Class A (Ballosteros-Weinstein ^12^), Class B1 and B2 (Wootten ^25^), Class C (Pin ^26^), Class D (fungal ^27^) and Class F (Wang ^28^). For G protein per-residue frustration density, Common G protein Numbering (CGN) was employed for alignment.

The mean residue frustration density 𝜂*_i_*^𝑋^ representing the frustration of a GPCR or G protein residue across *s* PDBs, was computed as:

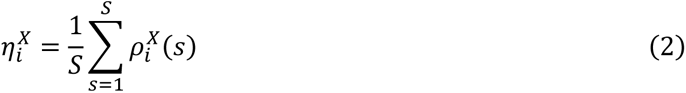

where the per-residue frustration density 𝜌*_i_*^𝑋^ is averaged across *S* structures. For GPCR calculation, per-residue frustration densities for each generic GPCR number were extracted from individual PDBs, and their mean value was calculated to represent the mean residue frustration density for that generic GPCR number. For G protein calculation, per-residue frustration densities for each CGN were extracted from individual structures, and their mean value was calculated to represent the mean residue frustration for that CGN.

Quartile analysis was performed for each GPCR class, and generic GPCR residues in the top quartile for high and minimal frustration were plotted onto representative structures in PYMOL. Quartile analysis was done on GPCR-Gα complex, Gα-Gβγ trimer, and Gα monomer groups separately, and CGN positions in the top quartile were plotted onto the representative structures in PYMOL.

The mean region frustration density 𝜂*_R_*^𝑋^, representing the frustration of a GPCR structural region 𝑅 across 𝑆 structures, was computed as:

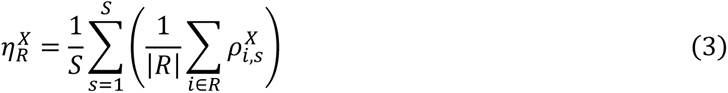

where we compute the mean frustration density for a region 𝑅 in each structure *S*, denoted as η*_R,s_*^𝑋^, by averaging the frustration densities ρ*_i,s_*^𝑋^ of all residues *i* in the region 𝑅. Here, |𝑅| represents the total number of residues in region *R*. We then compute 𝜂*_R_*^𝑋^ by averaging η*_R,s_*^𝑋^ over the available 𝑆 structures. A graphical representation of the schematics is depicted in Figure S1.

### Common highly frustrated residues in different Gα states

The top quartile of most frustrated CGN positions were collected from GPCR-Gα complex, Gα-Gβγ trimer, and Gα monomer groups. Only G_I-1_ and G_S_ subtypes have trimer structures, and therefore we only included structures of these two subtypes. CGN positions from HN helix and AHD regions were removed from the pool, because most GPCR-Gα complex structures do not have AHD regions, and most Gα monomer structures do not have HN helix. The Venn diagram was plotted using the Python matplotlib_venn package. Then, common highly frustrated GCN positions were grouped by the structural regions. The radial plots were created using Python Matplotlib.pyplot package using polar coordinates. Results from contact analysis were incorporated to plot the contacting region outer rings. If one GCN position from a structural region is in an interface, then the region is considered part of the interface.

### Allosteric modulator binding pocket identification and clustering

The structures of Class A GPCRs containing resolved PAM or NAM were collected and fed into Python environment via MDAnalysis package. For each structure, the coordinates of allosteric modulator heavy atoms were extracted, and the geometric center of the modulator was calculated using NumPy. The geometric centers of all available PAMs and NAMs were clustered separately using K-mean clustering in the SciPy package. The optimal number of clusters fed into K-means algorithm was determined by minimizing within-cluster sum of squares.

### Defining interface residues in apo structures

Bound structures of the same GPCR and same state (active: agonist-bound, inactive: antagonist-bound) were used for defining interface residues in apo structures. Due to the small size of apo structures, we used the interface residue list from each bound structure to calculate CDF in an apo structure.

### Engineered complex structure information

The three-dimensional structures of molecular glue degraders included cereblon, VHL, and DDB1-based degraders. 29 structures of the molecular glue complexes were that of DDB1– CDK12–cyclin K complexes. In these complexes the degrader mediates the interaction between DDB1 and CDK12, while cyclin K (Protein of Interest, POI) is not in direct contact with the degrader. This contrasts with most other molecular glue complexes, where the POI directly interacts with the degrader molecule. Hence, we separated these DDB1-based molecular glue complexes from VHL- or cereblon-based molecular glue complexes for buried surface area analysis.

Mutational frustration analysis was performed using FrustratometerR on a dataset consisting of 36 PROTAC-mediated complexes and 40 molecular glue complexes. The residue-based frustration was compared to 675 GPCR-Gα complexes. The PPI frustration percentage is defined as the ratio of highly (or minimally) frustrated contacts to the total number of PPI contacts. The PPI contacts refer to the inter-protein pairs identified by the FrustratometerR.

### Buried surface area calculation

The buried surface area was computed as:

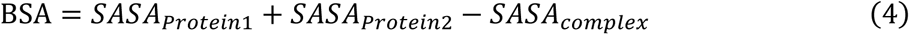

where SASA represents the Solvent Accessible Surface Area of the respective proteins and the complex.

### Interface residue polarization analysis

For amino acid polarization analysis, residues were classified as polar [R, N, D, Q, E, H, K, S, T, Y] or nonpolar [A, C, G, I, L, M, F, P, W, V] based on their side chain properties.

### Coevolutionary Analysis

We employed mutual information, an information-theoretic measure, to detect potential coevolutionary, structural, and functional connections between pairs of alignment sites. Given a multiple sequence alignment of 𝑛 sequences and 𝑝 positions, the coevolutionary score is derived from calculating the mutual information between positions according to:

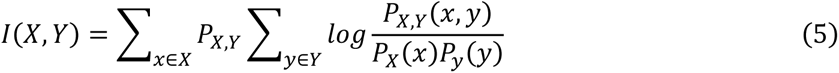

Here, 𝑃_𝑋_(𝑥) is the probability of observing an amino acid 𝑋 at position 𝑥 in the alignment, and 𝑃_𝑋,𝑌_(𝑥, 𝑦) is the joint probability of seeing an amino acid 𝑋 at position 𝑥 together with amino acid 𝑌 at position 𝑦. To highlight key interactions, the Z-score for positions 𝑥 and 𝑦 is then determined as:

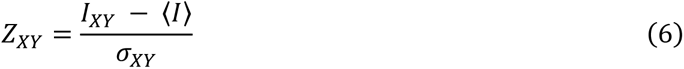

Once the coevolutionary scores for all position pairs have been computed using Equations (1) and (2), we condense these pairwise values into a single cumulative Z-score for each residue and rank them in descending order. Our premise is that residues exhibiting high Z-scores play important structural or functional roles, whereas those with lower Z-scores could be essential to receptor stability. We obtained the MSA of 287 human class A GPCR sequences for this step from GPCRdb (https://gpcrdb.org/).

### Conservation Analysis

For the conservation analysis, we used an MSA of 287 human class A GPCR sequences, obtained from the GPCR database (https://gpcrdb.org/). We computed a conservation score *C_i_* using Von Neumann entropy *S_i_* that incorporates chemical similarities among amino acids to yield a more robust metric of conservation across the alignment:

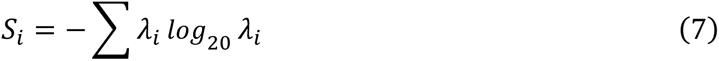

where 𝜆_𝑖_ is the eigenvalues of *ρ,* a density matrix based on the relative frequencies 𝑃(𝑌) of amino acids at each position in combination with a similarity matrix (BLOSUM50):

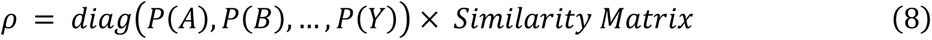

The final conservation score is found by subtracting the entropy from the maximum possible value, 1. Thus, 𝐶_𝑖_ = 1 − 𝑆_𝑖_.

## Acknowledgements

This work was done with grant funding from NIH R01-GM117923 and R35-GM156498 to N.V. and funding from Bristol-Myers-Squibb to N.V.

