## Supplementary figures for "Protein Frustration Reveals Active Sites in Co-Evolved GPCR:G Protein Complexes and in Engineered Targeted Degrader Complexes"

[3] Bristol-Myers-Squibb

[4] Department of Genetics, Cell Biology and Development, University of Minnesota, Minneapolis, MN, USA

### Supporting Information

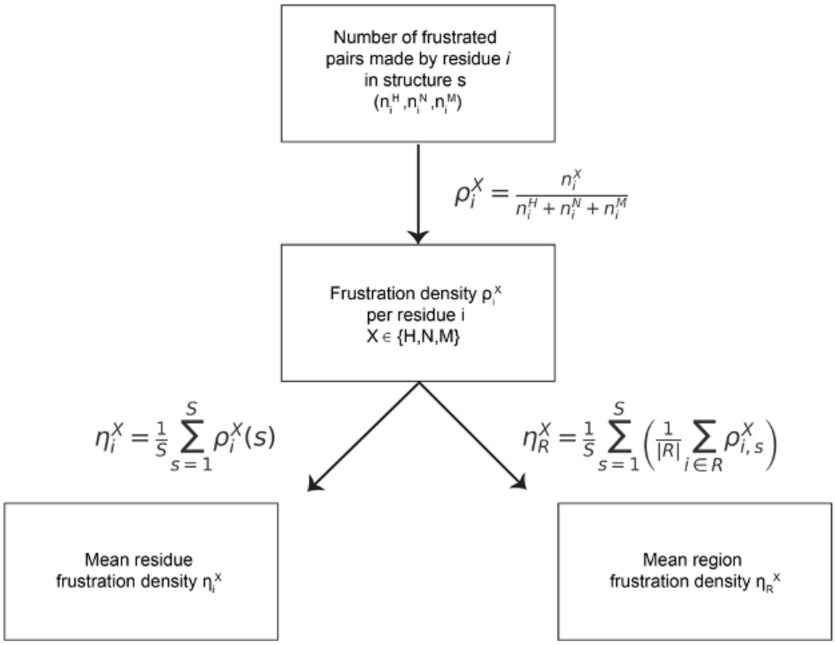


**Figure S1.** Workflow of calculating per-residue frustration density ($\rho_i^X$), mean residue frustration density ($\eta_i^X$), and mean region frustration density ($\eta_R^X$) from pairwise frustration across multiple PDB structures ($s$). Each residue $i$ interacts with $n_i^X$ residues within a 5 Å sphere, generating frustrated pairs categorized as highly frustrated ($n_i^H$), neutrally frustrated ($n_i^N$), or minimally frustrated ($n_i^M$). The frustration type is denoted by $X \in \{H, N, M\}$. $\rho_i^X$ is the per-residue frustration density for the residue $i$ and frustration type $X$. $\eta_i^X$ is the mean frustration density of the residue $i$ across $S$ structures. $\eta_R^X$ is the mean frustration density of a region $R$ across $S$ structures, where $|R|$ is the number of residues in the region $R$.

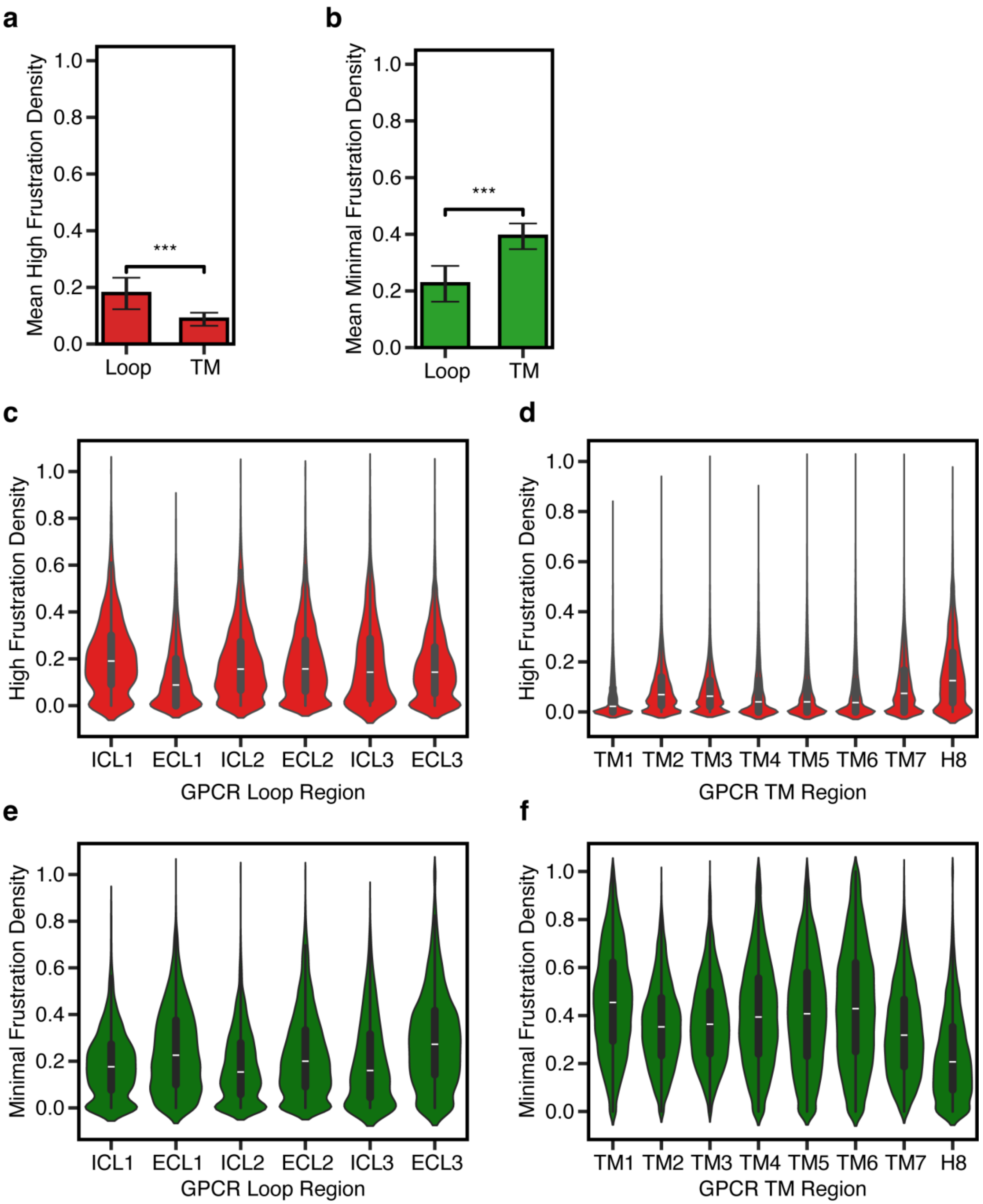


**Figure S2.** A, B) Bar plots comparing mean regional HF and MF density for residues located in GPCR loop regions (n = 1007) compared to those in TM regions (n = 1007). Data are shown as mean +/ SD. Comparison was performed using an independent t-test (two-tailed, unequal variances assumed). C, D) Violin plots comparing median regional HF frustration density ($\eta_R^X$) across different GPCR loop regions (C): ICL1 (n = 968), ECL1 (n = 989), ICL2 (n = 999), ECL2 (n = 1003), ICL3 (n = 543), and ECL3 (n = 972) and across different GPCR transmembrane regions (D): TM1 (n = 1006), TM2 (n = 1006), TM3 (n = 1007), TM4 (n = 1007), TM5 (n = 1006), TM6 (n = 996), TM7 (n = 999) and H8 (n = 948). E, F) Violin plots comparing median regional MF frustration density ($\eta_R^X$) across different GPCR loop regions (E): ICL1 (n = 968), ECL1 (n = 989), ICL2 (n = 999), ECL2 (n = 1003), ICL3 (n = 543), and ECL3 (n = 972) GPCR transmembrane regions (F): TM1 (n = 1006), TM2 (n = 1006), TM3 (n = 1007), TM4 (n = 1007), TM5 (n = 1006), TM6 (n = 996), TM7 (n = 999) and H8 (n = 948). Boxes represent the IQR with the median, and whiskers extend to the minimum and maximum values.

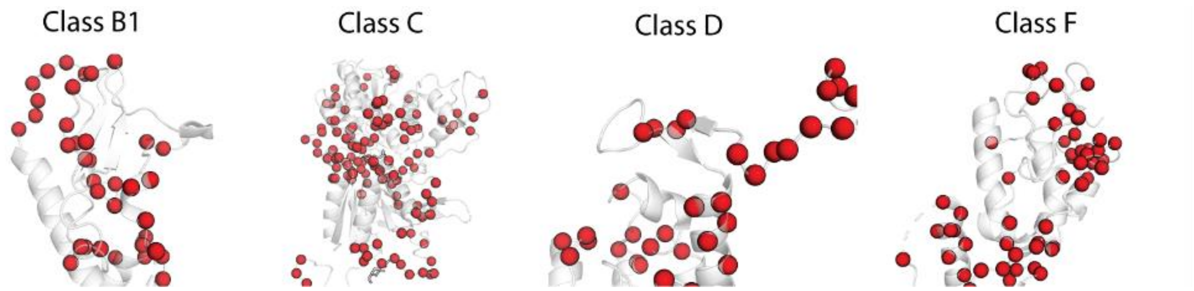


**Figure S3.** A) Highly frustrated residues in the Extracellular (EC) domain of Class B1 (PDB ID: 6X19), Class C (PDB ID: 7MTS), Class D (7AD3), and Class F (6XBK) GPCRs.

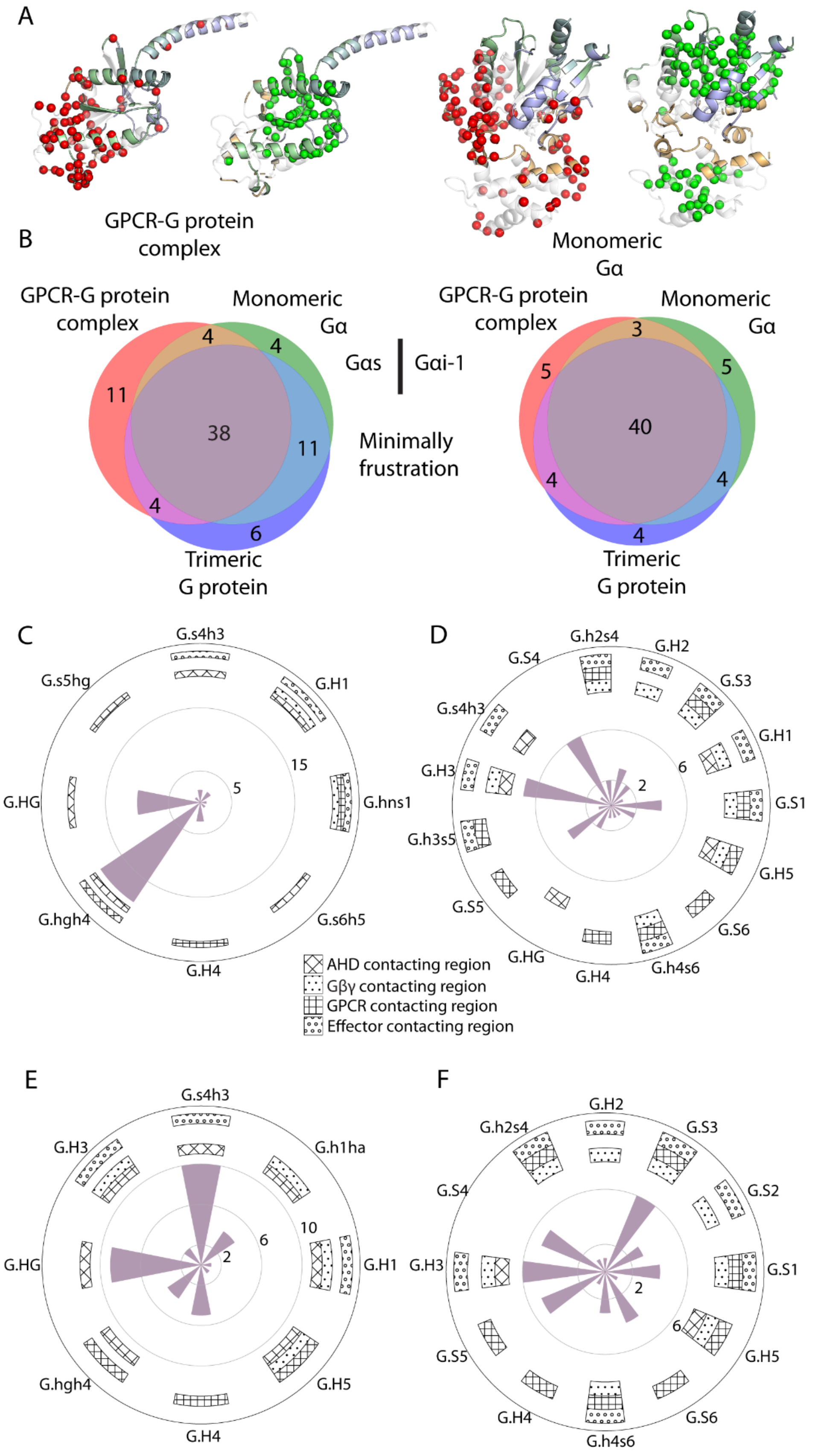
A
GPCR-G protein complex
Monomeric Gα
B
GPCR-G protein complex
Monomeric Gα
Gαs
Gαi-1
Minimally frustration
Trimeric G protein
11
4
4
38
11
4
6
5
3
5
40
4
4
4
C
D
E
F
AHD contacting region
Gβγ contacting region
GPCR contacting region
Effector contacting region
G.s4h3
G.H1
G.hns1
G.s6h5
G.H4
G.hgh4
G.HG
G.s5hg
G.h2s4
G.H2
G.S3
G.S1
G.H5
G.S6
G.h4s6
G.S5
G.h3s5
G.H3
G.S4
G.h1ha
G.S2

**Figure S4. Frustration of residues in Gα in GPCR-bound complex and GTP-bound monomer states.** A) High frustration (left) and minimal frustration residues (right) in Gα subunit in GPCR-bound Gα state (left) and GTP-bound Ga monomer state (right). B) Venn diagrams of the minimally frustrated residues (MFRs) in three conformational states (GDP-bound trimer state, GPCR-bound intermediate state, and GTP-bound Gα monomeric state) illustrating the common and distinct minimally frustrated residues in the three states in $G_S$ (left) and $G_{I\text{-}1}$ (right). C) Radial diagrams showing the substructure localization of commonly highly frustrated residues in $G\alpha_S$. D) Radial diagrams showing the location of the minimally frustrated residues in the structure of Gαs. E) Radial diagram showing the structural regions in which the commonly highly frustrated residues are located in $G\alpha_i$. F) Radial diagram showing the substructure localization of commonly minimally frustrated residues in $G\alpha_I$. The different patterns in the patches shown in C-F indicate whether this region in Gα forms the interface with an effector protein: diagonal cross – AHD interface; dotted – Gβγ interface; cross – GPCR interface; circle dots – other effectors.

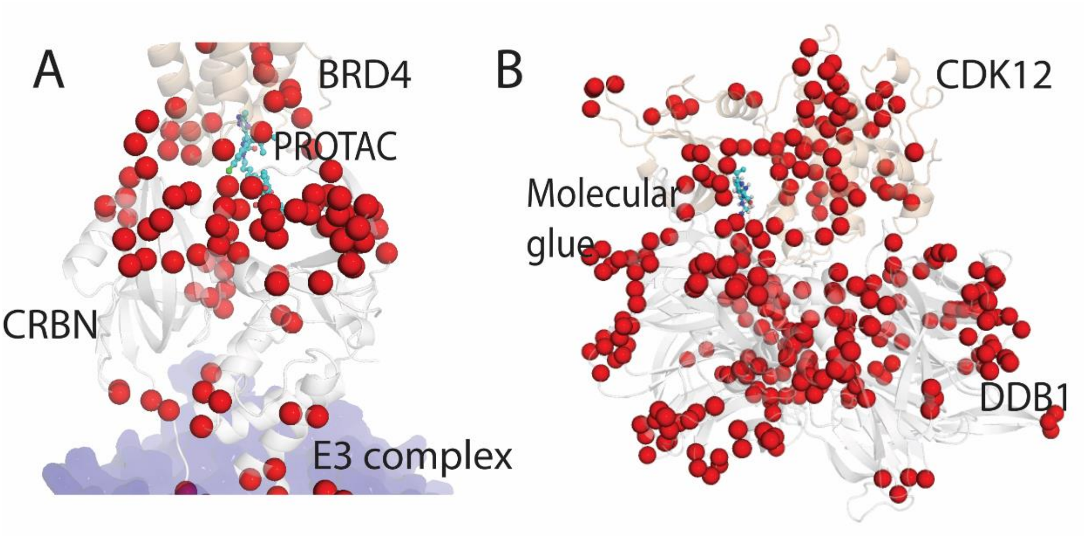


**Figure S5. Highly frustrated residues mapped on the PROTAC and molecular glue structure.** A) Highly frustrated residues mapped on DDB1-CRBN-PROTAC-BRD4 complex (PDB id: 6BOY). B) Highly frustrated residues mapped on DDB1-molecular glue-CDK12 complex (PDB id: 8BU1)

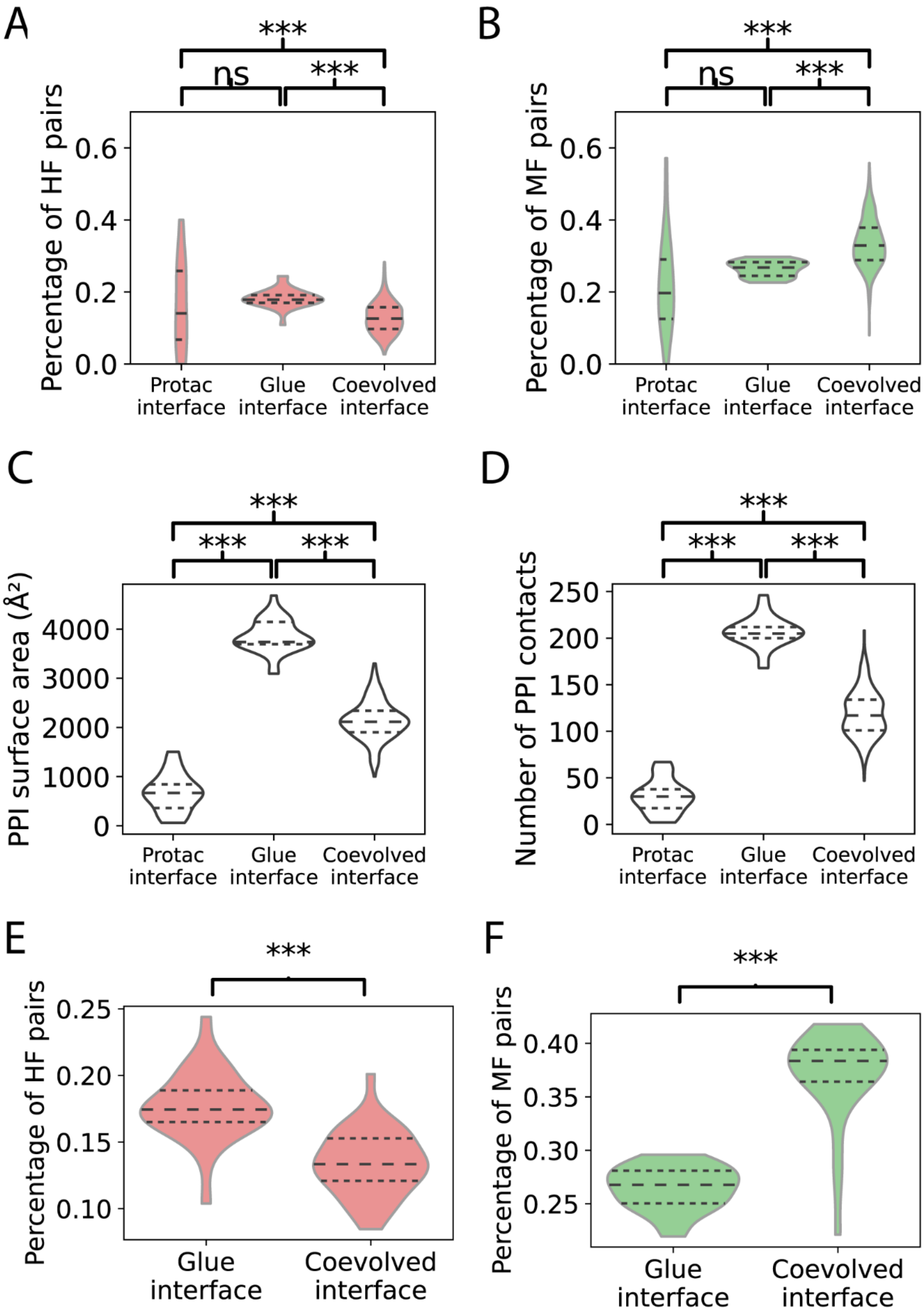

A
B
C
D
E
F
***
ns
Percentage of HF pairs
Percentage of MF pairs
PPI surface area (Å²)
Number of PPI contacts
Protac interface
Glue interface
Coevolved interface
0.0
0.2
0.4
0.6
0
1000
2000
3000
4000
50
100
150
200
250
0.10
0.15
0.20
0.25
0.30
0.35
0.40

**Figure S6.** The comparison of the molecular glue-mediated DDB1-CDK12 interface with the PROTAC-mediated and GPCR-G protein interface. A, B) High and minimal frustration level of residues in the protein-protein interface for PROTAC-mediated interface, molecular glue-mediated DDB1-CDK12 interface, and coevolved interface (GPCR-G protein interface). C) PPI surface area. D) Number of PPI contacting residue pairs. E, F) High and minimal PPI frustration level for molecular glue-mediated mediated DDB1-CDK12 complexes. The coevolved interface refers to the PPI between CDK12 and cyclin K.

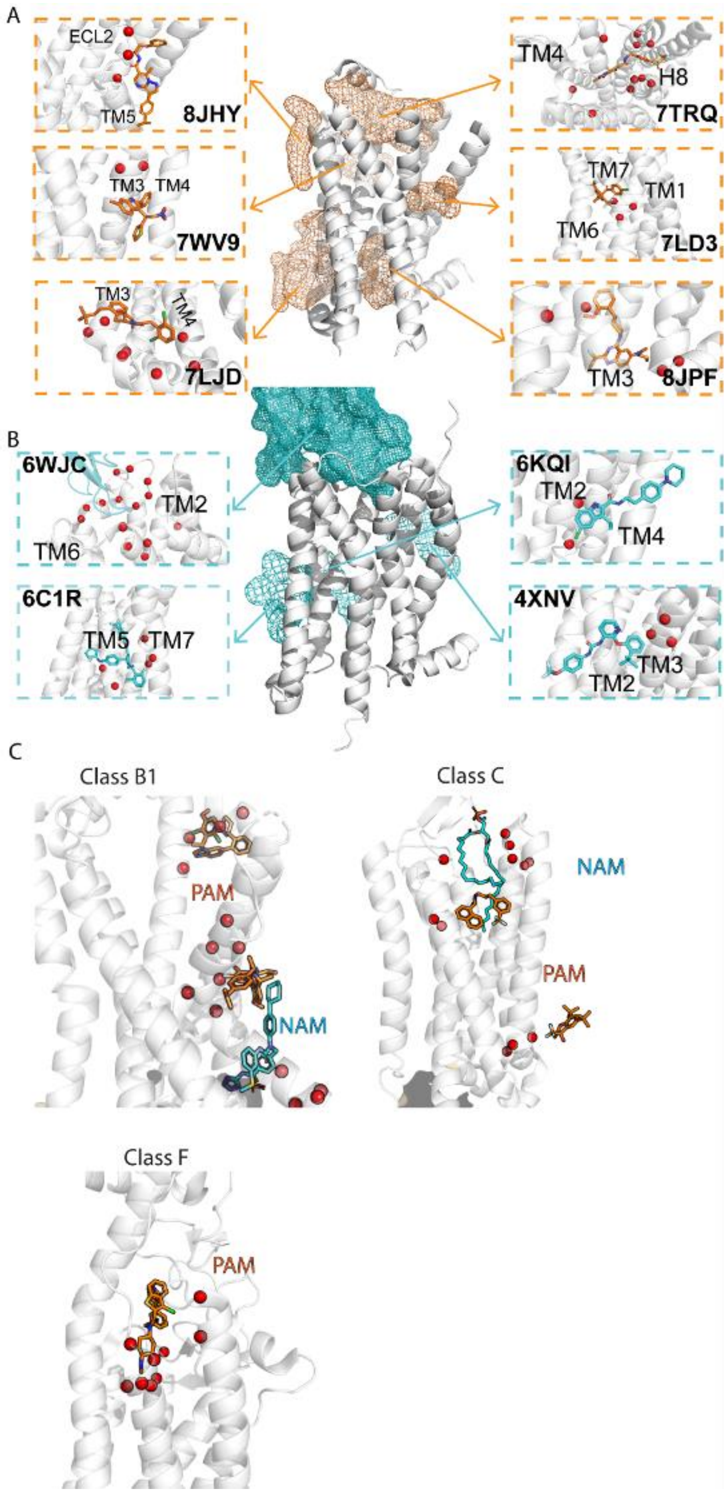


**Figure S7. Frustration profile of allosteric modulator binding sites.** A, B) The atoms density cloud of all known PAM (orange)/NAM (cyan). One PAM/NAM for each density cloud is shown as an example, with the highly frustrated PAM/NAM contacting residues shown as red spheres. C) Example of PAM/NAM and the highly frustrated contacting residues in their binding sites, for Class B1, C, and F GPCRs.
